# Safety Assessment of a Standardized Polyherbal Formulation (Glubloc™): Acute Toxicity Study and 28-Day Repeated Dose Toxicological Studies

**DOI:** 10.1101/2023.09.13.557515

**Authors:** Vinyas Mayasa, Pathuri Raghuveer, Pavan Kumar Chintamaneni, Chaitanya Chakravarthi Gali, Uttam Kumar Neerudu

## Abstract

The standardized extract of *Morus alba* and *Malus domestica Peel* is widely recognized as a dietary supplement with potential benefits for blood sugar management, cholesterol reduction, and weight management. This study aimed to evaluate the general toxicity of the extract. Toxicity studies were conducted by OECD guidelines 425 and 407 and Schedule Y of the Drugs and Cosmetics Act, which mandates the use of two species for such studies. A single dose of Glubloc^TM^ at a concentration of 2000 mg/kg body weight was administered to Sprague Dawley rats and Albino mice, and no fatalities were reported, indicating good tolerability. Additionally, repeated oral administration of Glubloc^TM^ to rats and rabbits for a maximum duration of 28 days, at doses up to 414.16 mg/kg body weight per day in rats and 207.08 mg/kg body weight per day in rabbits, did not result in significant changes in medical assessments, ocular evaluations, weight gains, feeding behavior, or pathological findings compared to the control group. Overall, this study suggests that Glubloc^TM^ is well-tolerated without inducing any toxicity. The extract’s No-Observed-Adverse-Effect Level (NOAEL) is considered to be 414.16 mg/kg per day when administered repeatedly for 28 days in rats and 207.08 mg/kg in rabbits.

## 1. Introduction

Blood glucose management is critical to maintaining optimal health, particularly in individuals with diabetes or those at risk of developing diabetes [1]. Over the past few years, there has been an evident increase in a growing interest in natural products as potential therapeutic agents for blood glucose management [2]. Among the vast array of botanicals investigated, *Malus domestica* (apple) and *Morus alba* (white mulberry) have emerged as promising candidates due to their complex phytochemical profiles and potential health benefits [3].

*Malus domestica*, a perennial fruit tree of the Rosaceae family, is widely cultivated worldwide for its delicious and nutritious fruit. The apple fruit is a rich source of various bioactive constituents, including polyphenols, flavonoids, and dietary fiber. The polyphenols found in *Malus domestica*, such as flavonols, catechins, and procyanidins, have undergone detailed exploration for their potent antioxidant properties and their potential impact on blood glucose management [4]. These polyphenols are believed to exert their beneficial effects through intricate mechanisms, including inhibition of carbohydrate-digesting enzymes, modulation of glucose transporters, and enhancement of insulin sensitivity. *Malus domestica* has been reported to possess remarkable anti-inflammatory, anti-cancer, and anti-ageing properties, which may further contribute to its potential role in blood glucose management [5–12].

On the other hand, *Morus alba*, a deciduous tree native to Asia, has been traditionally recognized for its medicinal properties. The leaves of *Morus alba* are known to contain a diverse array of bioactive constituents, including flavonoids, alkaloids, and imino sugars, including compounds like 2-O-α-D-galactopyranosyl deoxynojirimycin, fagomine and DNJ (1-deoxynojirimycin) [13]. It has been shown to exhibit inhibitory activity against alpha-glucosidase, an enzyme responsible for carbohydrate breakdown in the small intestine, resulting in a reduction of postprandial glucose levels [14]. Furthermore, *Morus alba* has also have been confirmed to exhibit potent antioxidant, anti-inflammatory, and anti-diabetic properties, which may augment its potential therapeutic effects in blood glucose management [13].

The bioactive constituents present in these botanicals, including polyphenols, flavonoids, alkaloids, and imino sugars, are thought to modulate various cellular pathways and molecular targets involved in glucose metabolism [15]. For instance, the inhibition of carbohydrate-digesting enzymes by polyphenols and DNJ may lead to a reduction in postprandial glucose levels, while the modulation of glucose transporters and enhancement of insulin sensitivity may improve glucose uptake and utilisation by cells [16]. Additionally, the anti-inflammatory and antioxidant properties of *Malus domestica* and *Morus alba* may help reduce oxidative stress and inflammation associated with insulin resistance and diabetes [16–17]. Recent research has also shed light on the potential influence of gut microbiota in blood glucose management and the interaction between botanicals and the microbial community residing in the gut, commonly known as gut microbiota, is increasingly recognised for its potential role in a critical role in glucose metabolism, and alterations in the gut microbiota composition have been associated with insulin resistance and diabetes [18–19]. *Malus domestica* and *Morus alba* have been shown to modulate the gut microbiota composition, promoting the growth of beneficial bacteria and inhibiting the proliferation of harmful bacteria [20–22]. This modulation of gut microbiota by these botanicals may contribute to their beneficial effects on blood glucose management through intricate mechanisms, such as short-chain fatty acid production, gut barrier function improvement, and modulation of gut hormone secretion [23–24].

Furthermore, the potential therapeutic applications of *Malus domestica* and *Morus alba* in blood glucose management extend beyond their direct effects on glucose metabolism [25]. These botanicals have been reported to possess anti-obesity properties, which may be beneficial in managing blood glucose levels, as obesity is a significant risk factor for diabetes [26]. Additionally, *Malus domestica* and *Morus alba* have been shown to possess cardiovascular health benefits, including cholesterol-lowering effects, anti-hypertensive effects, and improvement in endothelial function, which may indirectly contribute to blood glucose management through their effects on overall cardiovascular health [10,26,27]. The potential therapeutic applications of *Malus domestica* and *Morus alba* in blood glucose management have significant implications for public health. With the Mounting incidence of diabetes and the limitations of current treatment options, there is a pressing need to explore alternative approaches for blood glucose management [28]. The utilization of natural products, such as *Malus domestica* and *Morus alba*, as adjunctive or complementary therapies, may provide novel and promising avenues for improving blood glucose control and reducing the burden of diabetes. The intricate interplay and potential synergistic effects of combining *Malus domestica* and *Morus alba* have been the subject of recent investigations. The standardised extracts of *Malus domestica* and *Morus alba* have been formulated into proprietary blend, Glubloc™, Which have been demonstrated to exhibit enhanced effects on blood glucose management compared to individual extracts. These proprietary blends are patented and meticulously formulated to ensure consistent quality and potency, making them a promising avenue for further research and development.

## 2. Materials and methods

### 2.1. Test material

Glubloc™ is a standardised 2:3 proprietary blend of water extract of *Malus domestica* rind 4:1 extract with >40% polyphenolic flavonoids (Quercetin, procyanidins, catechins, phloridzin, phlorizin, epicatechin, rutin, and chlorogenic acid) and 5:1 *Morus alba* aqueous extract with >20% polyphenolic flavonoids and imino sugars (quercetin 3-rutinoside (rutin), quercetin 3-glucoside (isoquercetin), kaempferol 3-glucoside (astragalin), 1,5-dideoxy-1,5-imino-D-sorbitol hydrochloride (1-DNJ), fagomine, and 2-O-α-D-galactopyranosyl deoxynojirimycin). The final blend is a light brownish yellow-brown powder with a characteristic odor and taste and has been manufactured following strict cGMP practices.

### 2.2 Chemical Required

Sodium chloride, Potassium chloride, Sodium hydroxide, Hydrochloric acid, Ethanol, Acetone, Hematoxylin and Eosin, Erba kits were used for biochemical analysis.

### 2.3 Experimental Animals

We acquired adult Wistar rats (7-8 weeks old) and Swiss albino mice (6-8 weeks old) of both genders from Jeeva Life Sciences located in Hyderabad, India. These animals were free from pathogens. We obtained New Zealand white rabbits weighing 1.5 – 1.6 kg, aged 7-8 weeks, from Mahaveea Enterprises in Hyderabad, India. Before the commencement of the experiments, the animals were given a week to acclimate to the housing conditions, which included a temperature of 22 ± 3°C, relative humidity ranging from 40% to 70%, and a 12-hour light-dark cycle. The animals had unrestricted access to standard rodent chow and reverse osmosis-filtered water. All aspects of animal handling and investigations adhered to the guidelines set forth by the Committee for the Control and Supervision of Experiments on Animals (CCSEA) for animal care. The Institutional Animal Ethics Committee (IAEC) approved the study protocols at Jeeva Life Sciences in Telangana, India.

### 2.4. Acute Oral Toxicity Study

A study on acute oral toxicity was performed in both male and female wistar rats and Swiss albino mice, following the principles of Good Laboratory Practice (GLP) and by the OECD 425 guidelines for the testing of chemicals, as specified by the International [C(97)186/Final] legislation, by the Quality Assurance Unit of M/s. Jeeva Life Sciences, Hyderabad, Telangana, India with the study numbers JLS/021/PCT/PO/01/2023 & JLS/022/PCT/PO/01/2023 and the test item registration code provided as JLR/01/09/001/23.

A limit test was conducted using a sequential approach involving five wistar rats and albino mice with a 48-hour interval between each test. Glubloc™ was dissolved in distilled water, and a single oral dose of 2000 mg/kg was administered via oral gavage. After the administration, the animals were subjected to hourly monitoring for any signs of toxicity within the initial four hours, followed by daily observations for the subsequent 14 days. Comprehensive assessments were conducted to detect abnormalities in various functional parameters such as neuromuscular functions, excitability, sensory motor activities, and autonomic responses, including palpebral closure, urination, defecation, lacrimation, and more. Following the administration, the animals were monitored for any clinical signs of toxicity every hour for the first four hours and then every day for the next 14 days. All animals were observed for functional battery abnormalities, including neuromuscular excitability, sensory-motor activities, and autonomic parameters, including palpebral closure, urination, defecation, lacrimation, etc. Detailed clinical examination was conducted on days 1, 7, and 14 for all animals. After euthanasia on day 15, a thorough analysis of vital organs and tissues was conducted to identify any visible gross pathological changes [29,30].

### 2.5. Twenty-Eight-Day Repeated Dose Oral Toxicity Study

A study on repeated oral toxicity lasting for twenty-eight days was conducted (ethics committee approval numbers JLS/023/PCT/PO/01/2023 and JLS/024/PCT/PO/01/2023) was conducted in Wistar rats and New Zealand white rabbits (both sexes), following the OECD guidelines 407, in compliance with OECD principles of GLP for the testing of chemicals as specified by International [C(97)186/Final] legislation by the Quality Assurance Unit of M/s. Jeeva Life Sciences [31].

Wistar rats of both sexes (n = 48; 24 males and 24 females) were randomly allocated into four groups (n = 12; 6 males and 6 females/ group). Test supplement, Glubloc™, prepared in distilled water was administered daily for twenty-eight days in single oral doses of 103.54 mg/kg (low dose, G2), 207.08 mg/kg (mid dose, G3), or 414.16 mg/kg BW (high dose, G4). The vehicle control group (G1) received distilled water with no test item (0) through oral gavage daily for 28 days. The test item formulation was prepared freshly every day before dosing the animals; water is used as the diluent to get the desired concentration of 103.54 mg/kg/5ml, 207.08 mg/kg/5ml and 414.16 mg/kg/5ml for G2, G3 and G4 of the test item.

New Zealand white rabbits of both sexes (n = 24; 12 males and 12 females) were randomly allocated into four groups (n = 6; 3 males and 3 females/ group). Test supplement, Glubloc™, prepared in distilled water was administered daily for twenty-eight days in single oral doses of 51.77 mg/kg (low dose, G2), 103.54 mg/kg (mid dose, G3), or 207.08 mg/kg BW (high dose, G4). The vehicle control group (G1) received distilled water with no test item (0) through oral gavage daily for 28 days. The test item formulation was prepared freshly every day before dosing the animals, water was used as the diluent to get the desired concentration of 51.77 mg/kg/5ml, 103.54 mg/kg/5ml and 207.08 mg/kg/5ml for G2, G3 and G4 of the test item.

During the necropsy, the animals underwent an overnight fasting period, and blood samples were obtained through retro-orbital puncture (while under isoflurane anaesthesia) to perform haematology and clinical chemistry analyses. Hematobiochemical parameters were assessed using the ADVIA 2120 haematology analyser (Siemens Healthcare Private Limited, Munich, Germany) and a clinical chemistry analyzer (ILab Aries, Bergamo, Italy). Various hemotological and biochemical parameters were analysed. Following euthanasia using CO_2_, the vital organs of the animals were gathered for both macroscopic and microscopic pathological examinations. The collected organs included the liver, kidneys, heart, lungs, brain, spleen, adrenal glands, thymus, testes, epididymis, seminal vesicles, ovaries, and uterus. The organs were weighed using a high-precision electronic balance with an accuracy of 0.01 g (Mettler Toledo, Columbus, OH). To conduct histopathology examinations, the organs were immersed in 10% neutral buffered formalin for 48 hours for fixation. Subsequently, the organs embedded in paraffin were sectioned into slices with a thickness of 5 μm using a rotary microtome. The tissue sections underwent processing in alcohol of varying concentrations and were then stained with hematoxylin and eosin. Subsequently, the stained tissue sections were examined using a light microscope.

### 2.6. Statistical Analyses

The data were presented as the mean ± standard deviation (SD). Comparative analyses between different groups were conducted using one-way analysis of variance (ANOVA) followed by Dunnett’s post hoc test. All comparisons were assessed with a 95% confidence level, and statistical significance was determined by P values below 0.05.

## 3. Results

### 3.1. Acute toxicity study

The administration of a single oral dose of Glubloc™ did not cause functional disturbance or death in Wistar rats and Swiss Albino mice. No morbidity or mortality were observed in any of the groups following a single-dose oral administration of 2000 mg/kg BW Glubloc™ in the Wistar rats and Swiss Albino mice of both sexes. The animals were active and did not show any visible signs of toxicity. None of the groups showed any abnormal changes in their body weights, feeding patterns, or water consumption during the post-dose 14-day observation period. No abnormalities were detected in the functional battery or gross pathological examination of animals at the end of the study period. Mean summary body weights of the rats and mice were shown in table −2 and 3 for both s. No statistical significance in the bodyweights and food consumption was observed. Together, these observations indicate that the median oral lethal dose (LD_50_) of Glubloc™ is at least 2000 mg/kg BW in Wistar rats and Swiss Albino mice (both sexes)

**Table 1.** Specifications of Glubloc™. (Supplementary Material)

| Parameter | Specification | Assay Method |
| --- | --- | --- |
| Description | Brownish-yellow to brown powder | Organoleptic |
| Odor | Characteristic | Organoleptic |
| Taste | Characteristic | Organoleptic |
| Loss on drying | <5% | USP <731> |
| Solubility (1 g/300 ml water) | NLT 80% w/w | USP General Chapter |
| pH (1% w/v aq. solution) | 4.0 – 7.0 | pH meter |
| Particle size | 100% thru 80 mesh | USP <786> |
| Tap density | 0.4 – 0.95 g/cc | USP <616> |
| Bulk density | 0.3 – 0.8 g/cc | USP <616> |
| **Chemical assay** |  |  |
| Polyphenols | NLT 20% w/w dry extract | HPLC |
| Flavonoids | NLT 10% w/w dry extract | HPLC |
| Heavy metals | NMT 10 ppm | AAS |
| Lead | NMT 3.0 ppm | AAS |
| Mercury | NMT 0.1 ppm | AAS |
| Cadmium | NMT 1.0 ppm | AAS |
| Arsenic | NMT 2.0 ppm | AAS |
| **Microbiological assays** |  |  |
| Total plate count (CFU/g) | Less than 10,000 | USP <2021> |
| Yeast and mould (CFU/g) | Less than 1000 | USP <2021> |
| E. coli (CFU/g) | Negative | USP <2022> |
| Salmonella (CFU/375 g) | Negative | USP <2022> |
| S. aureus (CFU/25 g) | Negative | USP <2022> |
| **Pesticide assays** |  | USP <561> |
| Organochlorine | Negative |  |
| Organophosphorus | Negative |  |
| Organonitrogen | Negative |  |
| N-Methyl carbamates | Negative |  |
| Aflatoxins | Negative |  |

AAS = Atomic Absorption Spectroscopy; CFU = Colony Forming Units; HPLC = High-Performance Liquid Chromatography; USP = United States Pharmacopeia.

**Table 2:** Summary of Mean Body Weight (g) - rats.

| Groups (Sex) | Treatment | Mean Body weight (g) |  |  |
| --- | --- | --- | --- | --- |
|  |  | Day 1 | Day 7 | Day 14 |
| **G1-Male** | **Vehicle control** | 185.1±3.06 | 198.6±3.67<br>(6.56%) | 210.5±3.26<br>( 11.9%) |
| **G2- Male** | **Glubloc™**<br>**(2000mg/Kg)** | 183.6±4.00 | 207.08.3±4.75<br>(8.5%) | 215.2±2.75<br>( 14.8%) |
| **G1-Female** | **Vehicle control** | 183.8±2.34 | 198.5±4.02<br>(7.57%) | 213.4±2.56<br>( 15.02%) |
| **G2-Female** | **Glubloc™**<br>**(2000mg/Kg)** | 188.9±1.16 | 207.08.2±1.37<br>(6%) | 215.8±2.42<br>(12.55%) |

The data is reported as mean ± SD for a sample size of 6 (n=6). The symbols \* and # indicate statistical significance with p-values less than 0.05 compared to G1 for each symbol, respectively.

**Table 3:** Summary of Mean Body Weight (g) - Swiss Albino mice.

| Groups | Treatment | Mean Body weight (g) |  |  |
| --- | --- | --- | --- | --- |
|  |  | Day 1 | Day 7 | Day 14 |
| **G1-Male** | **Vehicle control** | 18.16±2.06 | 22.02±2.67<br>(18.18%) | 25.17±2.26<br>(24.28%) |
| **G2- Male** | **Glubloc™**<br>**(2000 mg/Kg)** | 19.66±2.03 | 23.04±2.75<br>(17.39%) | 25.16±2.75<br>(24%) |
| **G1-Female** | **Vehicle control** | 20.26±2.14 | 21.15±2.07<br>(20.04%) | 23.17±2.22<br>(13.04%) |
| **G2-Female** | **Glubloc™**<br>**(2000mg/Kg)** | 20.66±2.03 | 22.04±2.18<br>(9.09%) | 24.16±2.75<br>(16.66%) |

The data is reported as mean ± SD for a sample size of 5 (n=6). The symbols \* and # indicate statistical significance with p-values less than 0.05 compared to G1 for each symbol, respectively.

### 3.2. Repeated dose oral toxicity studies

Twenty-eight-day repeated dose oral toxicity study with Glubloc™ was nontoxic and yielded no unwarranted effects in New Zealand white rabbits and Wistar rats.

Throughout the study, there were no recorded mortality occurrences in either the control or treated groups. Regular and thorough clinical and functional observations of the animals uncovered no significant toxicological abnormalities. No significant changes related to the test item were observed in the body weight (refer to Supplemental Figures 1a and 1b), body weight gain, or daily food consumption of male or female New Zealand white rabbits and Wistar rats at any of the illustrated dose levels in graphical images 1 and 2, as well as tables 4 and 5. No alterations were noted in physical examinations. No statistically significant changes were noted in hematology and clinical chemistry parameters in both sexes compared to control animals (see Tables 8 and 9). Table-4 displays the impact of Glubloc^TM^ on the percentage change in body weight, revealing a significant increase in body weight across all dose levels in comparison to the respective initial values. These findings indicate that the test drug does not harm animals’ health status, as evidenced by the weight gain observed during the experimental period.

**Table 4:** Summary of Food Consumption/Animal/Day (g) – (Swiss Albino mice)

| Groups | Treatment | Mean Food Consumption (g) |  |  |
| --- | --- | --- | --- | --- |
|  |  | Day 1 | Day 7 | Day 14 |
| G1-Male | Vehicle control | 2.24 | 2.79<br>(19.71%) | 2.34<br>(4.27%) |
| G2- Male | Glubloc™<br>(2000 mg/Kg) | 2.76 | 2.18<br>(26.60 %) | 2.97<br>(7.07%) |
| G1-Female | Vehicle control | 2.27 | 2.56<br>(11.32%) | 2.16<br>(5.09%) |
| G2-Female | Glubloc™<br>(2000 mg/Kg) | 2.14 | 2.36<br>(9.32%) | 2.06<br>(3.88%) |

The data is reported as mean ± SD for a sample size of 5 (n=6). The symbols \* and # indicate statistical significance with p-values less than 0.05 compared to G1 for each symbol, respectively.

**Table 5:** Summary of Mean Body Weight – Wistar Rats.

| Groups<br>Treatment | Mean Body weight (g)-Males |  |  |  |  | Mean Body weight (g)- Females |  |  |  |  |
| --- | --- | --- | --- | --- | --- | --- | --- | --- | --- | --- |
|  | Day 1 | Day 8 | Day 15 | Day 22 | Day 28 | Day 1 | Day 8 | Day 15 | Day 22 | Day 28 |
| G1 | 162.16±<br>1.06 | 182±<br>1.67 | 195.33±<br>3.26 | 214.66±<br>1.63 | 229±2.0 | 163.5±<br>2.34 | 177.16±<br>4.02 | 195.83±<br>2.56 | 214.5±<br>1.04 | 224<br>2.44 |
| G2 | 163.66 ±<br>1.03 | 180.83<br>±<br>2.75 | 192.83±<br>2.75 | 214.16±<br>1.47 | 225.83±<br>1.6 | 152.27±<br>1.16 | 164.51±<br>1.37 | 180.24±<br>2.42 | 194.89±<br>2.06 | 207.24<br>2.04 |
| G3 | 162.33 ±<br>1.21 | 175±<br>2.36 | 189.16±<br>2.31 | 212.33 ±<br>1.21 | 224.16±<br>2.31 | 162±<br>1.54 | 174.16±<br>0.75 | 192±<br>2.36 | 206.5±<br>2.58 | 221.33<br>4.92 |
| G4 | 163.33±<br>1.5 | 178.33±<br>1.86 | 192.33±<br>2.16 | 213.16±<br>1.72 | 225±<br>2.09 | 163.16±<br>0.68 | 182±<br>1.15 | 193.16±<br>1.46 | 205±<br>2.08 | 216.16<br>2.11 |

The data is reported as mean ± SD (n=6). The symbols \* and # indicate statistical significance with p-values less than 0.05 compared to G1 for each symbol, respectively.

**Table 6:** Summary of Mean Body Weight (g) - Rabbits.

| Treatment | Body weight (kg) (n=3; Males) |  |  |  |  | Bodyweight (Kg) (n=3; Females) |  |  |  |  |
| --- | --- | --- | --- | --- | --- | --- | --- | --- | --- | --- |
|  | Day 1 | Day 8 | Day 15 | Day 22 | Day 28 | Day 1 | Day 8 | Day 15 | Day 22 | Day 28 |
| G1 | 1.52±<br>0.21 | 1.59±<br>0.22 | 1.67±<br>0.31 | 1.8±<br>0.21 | 1.94±<br>0.21 | 1.53±<br>0.22 | 1.66±<br>0.14 | 1.84±<br>0.11 | 1.89±<br>0.04 | 1.99±<br>0.21 |
| G2 | 1.53±<br>0.31 | 1.62±<br>0.21 | 1.74±<br>0.31 | 1.85±<br>0.13 | 2.01±<br>0.35 | 1.53±<br>0.32 | 1.65±<br>0.11 | 1.76±<br>0.21 | 1.86±<br>0.32 | 1.97±<br>0.21 |
| **G3** | 1.52±<br>0.31 | 1.63±<br>0.31 | 1.74±<br>0.32 | 1.84±<br>0.35 | 2.03±<br>0.34 | 1.53±<br>0.11 | 1.62±<br>0.25 | 1.73±<br>0.31 | 1.85±<br>0.31 | 1.97±<br>0.22 |
| **G4** | 1.53±<br>0.5 | 1.66±<br>0.25 | 1.76±<br>0.32 | 1.88±<br>0.11 | 2.04±<br>0.31 | 1.54±<br>0.15 | 1.65±<br>0.21 | 1.76±<br>0.25 | 1.86±<br>0.31 | 1.96±<br>0.21 |

The data is reported as mean ± SD (n=3). The symbols \* and # indicate statistical significance with p-values less than 0.05 compared to G1 for each symbol, respectively.

**TABLE 7:** Summary of food consumption/animal/day (G)

| Groups | Treatment | Males (n = 3; Mean±SD) |  |  |  |  |  |  |
| --- | --- | --- | --- | --- | --- | --- | --- | --- |
|  |  | Day 4 | Day 8 | Day 12 | Day 16 | Day 20 | Day 24 | Day 28 |
| **G1** | **(Vehicle control)** | 163.00<br>±0.40 | 177.06<br>±1.82 | 191.08±<br>0.56 | 201.52<br>±2.05 | 211.80<br>±0.46 | 221.35<br>±1.07 | 228.34<br>±1.63 |
| **G2** | **Glubloc™**<br>**(51.77**<br>**mg/kg )** | 172.00<br>±0.57 | 172.58<br>±0.81 | 190.03<br>±2.28 | 201.21<br>±2.28 | 210.18<br>±1.34 | 217.91<br>±1.8 | 228.43<br>±0.69 |
| **G2** | **Glubloc™**<br>**(103.54**<br>**mg/kg )** | 163.15<br>±0.60 | 172.93<br>±0.66 | 185.75<br>±1.07 | 197.12<br>±1.10 | 207.11<br>±1.56 | 217.20<br>±0.59 | 226.67<br>±0.91 |
| **G4** | **Glubloc™**<br>**(207.08**<br>**mg/kg )** | 162.67±1.24 | 172.49<br>±0.78 | 184.6<br>±0.90 | 196.94<br>±0.66 | 206.33<br>±0.35 | 216.65<br>±0.68 | 225.83<br>±1.06 |

| Groups | Treatment | Females (n = 3; Mean±SD) |  |  |  |  |  |  |
| --- | --- | --- | --- | --- | --- | --- | --- | --- |
|  |  | Day 4 | Day 8 | Day 12 | Day 16 | Day 20 | Day 24 | Day 28 |
| **G1** | **(Vehicle control)** | 163.69<br>±1.12 | 174.33<br>±0.86 | 185.98<br>±1.34 | 196.47<br>±1.03 | 205.54<br>±1.14 | 214.39<br>±0.98 | 225.41<br>±0.93 |
| **G2** | **Glubloc™**<br>**(51.77**<br>**mg/kg )** | 162.19<br>±1.27 | 172.8<br>±1.01 | 184.58<br>±1.09 | 195.51<br>±0.76 | 205.16<br>±0.57 | 214.88<br>±1.47 | 224.32<br>±1.44 |
| **G2** | **Glubloc™**<br>**(103.54**<br>**mg/kg )** | 162.64<br>±1.22 | 172.62<br>±0.98 | 182.81<br>±1.72 | 195.21<br>±1.34 | 204.65<br>±1.57 | 213.66<br>±1.48 | 224.52<br>±1.05 |
| **G4** | **Glubloc™**<br>**(207.08**<br>**mg/kg )** | 162.94<br>±1.25 | 172.32<br>±2.04 | 182.82<br>±1.12 | 193.74<br>±1.4 | 203.58<br>±1.91 | 214.65<br>±0.38 | 223.34<br>±2.14 |

**TABLE 8.** Summary of hematology parameters in Wistar Rats.

| Males (n = 6; Mean±SD) |  |  |  |  |  | Females (n = 6; Mean±SD) |  |  |  |
| --- | --- | --- | --- | --- | --- | --- | --- | --- | --- |
| Parameter | Units | G1 | G2 | G3 | G4 | G1 | G2 | G3 | G4 |
| Total WBC | (10<sup>3</sup>/μl) | 12.89<br>±0.53 | 12.58<br>±0.41 | 11.7<br>±0.49 | 12.18<br>±0.71 | 12.53<br>±0.23 | 12.46<br>±0.43 | 12.81<br>±0.21 | 13.29<br>±0.31 |
| Total RBC | (10<sup>6</sup>/μL) | 9.19<br>±0.12 | 8.97<br>±0.21 | 8.04<br>±0.48 | 7.78<br>±0.31 | 9.35<br>±0.16 | 9.47<br>±0.16 | 8.57<br>±0.2 | 7.91<br>±0.44 |
| HB | (%) | 14.84<br>±0.44 | 14.72<br>±0.36 | 15.02<br>±0.61 | 14.14<br>±0.34 | 14.76<br>±0.67 | 12.21<br>±0.31 | 14.85<br>±0.43 | 14<br>±0.41 |
| PCV | (%) | 45.88<br>±1.25 | 42.47<br>±1.73 | 43.24<br>±1.49 | 43.07<br>±0.89 | 47.4<br>±1.86 | 47.19<br>±1.34 | 45.42<br>±1.2 | 45.24<br>±0.63 |
| MCV | (fl) | 55.86<br>±0.56 | 54.78<br>±1.57 | 52.96<br>±1.18 | 54.19<br>±1.2 | 55.38<br>±0.91 | 55.93<br>±1.17 | 55.77<br>±1.42 | 54.76<br>±1.32 |
| MCH | (pg) | 18.39<br>±0.19 | 17.91<br>±0.59 | 17.93<br>±0.43 | 17.88<br>±0.37 | 18.71<br>±0.56 | 18<br>±0.41 | 17.97<br>±0.26 | 17.89<br>±0.54 |
| MCHC | (%) | 32.85<br>±0.88 | 32.24<br>±0.93 | 33.32<br>±1.16 | 32.26<br>±1.26 | 32.47<br>±0.83 | 32.8<br>±1.12 | 32.9<br>±0.57 | 32.93<br>±0.31 |
| Platelets | (10<sup>3</sup>/μl) | 842<br>±9.93 | 838.5<br>±7.28 | 789.5<br>±10.98 | 796.6<br>±4.45 | 852<br>±5.44 | 847<br>±4.32 | 790.66<br>±6.77 | 793.66<br>±5.35 |
| Neutrophils | (%) | 21.74<br>±1.27 | 21.86<br>±1.09 | 22.86<br>±0.79 | 23.15<br>±0.7 | 22.08<br>±0.29 | 21.41<br>±0.79 | 22.11<br>±0.6 | 22.21<br>±0.75 |
| Lymphocytes | (%) | 77.33<br>±1.61 | 76.66<br>±1.77 | 77.69<br>±1.43 | 80.32<br>±1.15 | 79.3<br>±0.98 | 79<br>±0.91 | 79.13<br>±0.4 | 80.8<br>±0.79 |
| Eosinophils | (%) | 1.63<br>±0.1 | 1.6<br>±0.04 | 1.62<br>±0.04 | 1.63<br>±0.06 | 1.4<br>±0.11 | 1.98<br>±0.03 | 1.5<br>±0.05 | 1.57<br>±0.07 |
| Basophils | (%) | 0.003 | 0.002 | 0.003 | 0.001 | 0.005 | 0.004 | 0.004 | 0.003 |
| Monocytes | (%) | 2.62<br>±0.12 | 2.6<br>±0.007 | 2.45<br>±0.16 | 2.6<br>±0.17 | 2.59<br>±0.09 | 2.56<br>±0.07 | 2.47<br>±0.17 | 2.56<br>±0.06 |

The data is reported as mean ± SD for a sample size of 6 (n=6). The symbols \* and # indicate statistical significance with p-values less than 0.05 compared to G1 for each symbol, respectively.

**TABLE 8:** Summary of clinical chemistry parameters in New Zealand White Rabbits.

|  |  | Males (n = 3; Mean±SD) |  |  |  | Females (n = 3; Mean±SD) |  |  |  |
| --- | --- | --- | --- | --- | --- | --- | --- | --- | --- |
| Parameter | Units | G1 | G2 | G3 | G4 | G1 | G2 | G3 | G4 |
| Total WBC | (10<sup>3</sup>/μl) | 15.23<br>±0.51 | 15.75<br>±0.32 | 16.74±<br>0.48 | 17.03<br>±0.14 | 17.18<br>±0.11 | 17.44<br>±0.17 | 17.63<br>±0.12 | 18.15<br>±0.13 |
| Total RBC | (10<sup>6</sup>/μL) | 7.37<br>±0.17 | 7.32±<br>0.3 | 8.11±<br>0.13 | 8.28<br>±0.69 | 7.71<br>±0.23 | 7.43<br>±0.16 | 7.49<br>±0.1 | 7.58<br>±0.03 |
| HB | (%) | 15.46<br>±0.55 | 15.27±<br>0.04 | 15.53±<br>0.24 | 16.13<br>±0.14 | 15.02<br>±0.03 | 15.01<br>±0.03 | 14.99<br>±0.05 | 15.03<br>±0.16 |
| PCV | (%) | 34.83<br>±1.73 | 36.37±<br>1.23 | 35.72±<br>0.49 | 36.47<br>±0.43 | 36.14<br>±0.93 | 37.19<br>±0.64 | 35.73<br>±0.41 | 36.42<br>±0.89 |
| MCV | (fl) | 77.16<br>±0.98 | 77.58±<br>1.25 | 77.85±<br>0.29 | 78.46<br>±0.39 | 78.80<br>±0.64 | 78.83<br>±0.61 | 77.84<br>±1.57 | 77.99<br>±0.46 |
| MCH | (pg) | 25.92<br>±1.3 | 24.02±<br>1.23 | 24.62±<br>0.48 | 24.79<br>±1.13 | 25.63<br>±1.94 | 25.45<br>±0.59 | 26.67<br>±0.45 | 27.43<br>±1.35 |
| MCHC | (%) | 35.06<br>±1.2 | 34.87±<br>1.51 | 34.58±<br>0.8 | 34.85<br>±1.54 | 35.41<br>±0.79 | 36.46<br>±0.78 | 36.73<br>±0.68 | 37.19<br>±1.31 |
| Platelets | (10<sup>3</sup>/μl) | 552±<br>12.71 | 564±<br>19.26 | 554.66±<br>19.29 | 561.00<br>±18.18 | 581.00<br>±3.39 | 596.00<br>±23 | 589.33<br>±2.08 | 585.66<br>±3.51 |
| Neutrophils | (%) | 23.47<br>±0.59 | 23.75±<br>1.11 | 23.77±<br>1.36 | 22.06±<br>0.66 | 25.89<br>±0.49 | 26.11<br>±0.63 | 25.98<br>±0.51 | 25.49<br>±1.25 |
| Lymphocytes | (%) | 74.76<br>±0.92 | 75.16±<br>1.15 | 75.24±<br>0.9 | 76.68±<br>1.65 | 76.32<br>±0.37 | 76.38<br>±1.29 | 76.9<br>±0.07 | 76.52<br>±0.43 |
| Eosinophils | (%) | 2.47±<br>0.08 | 2.47<br>±0.18 | 2.65±<br>0.32 | 2.65±<br>0.19 | 2.70<br>±0.1 | 2.73<br>±0.09 | 2.54<br>±0.11 | 2.58<br>±0.09 |
| Basophils | (%) | 2.79±<br>0.15 | 2.71<br>±0.15 | 2.57±<br>0.4 | 2.63±<br>0.55 | 3.31<br>±0.16 | 3.2<br>±0.08 | 3.26<br>±0.07 | 3.27<br>±0.16 |
| Monocytes | (%) | 2.93±<br>0.55 | 3.35<br>±0.17 | 3.26±<br>0.02 | 3.3±<br>0.2 | 3.79<br>±0.05 | 3.77<br>±0.18 | 3.76<br>±0.311 | 3.81<br>±0.31 |

The data is reported as mean ± SD for a sample size of 3 (n=3). The symbols \* and # indicate statistical significance with p-values less than 0.05 compared to G1 for each symbol, respectively.

**TABLE 9:**
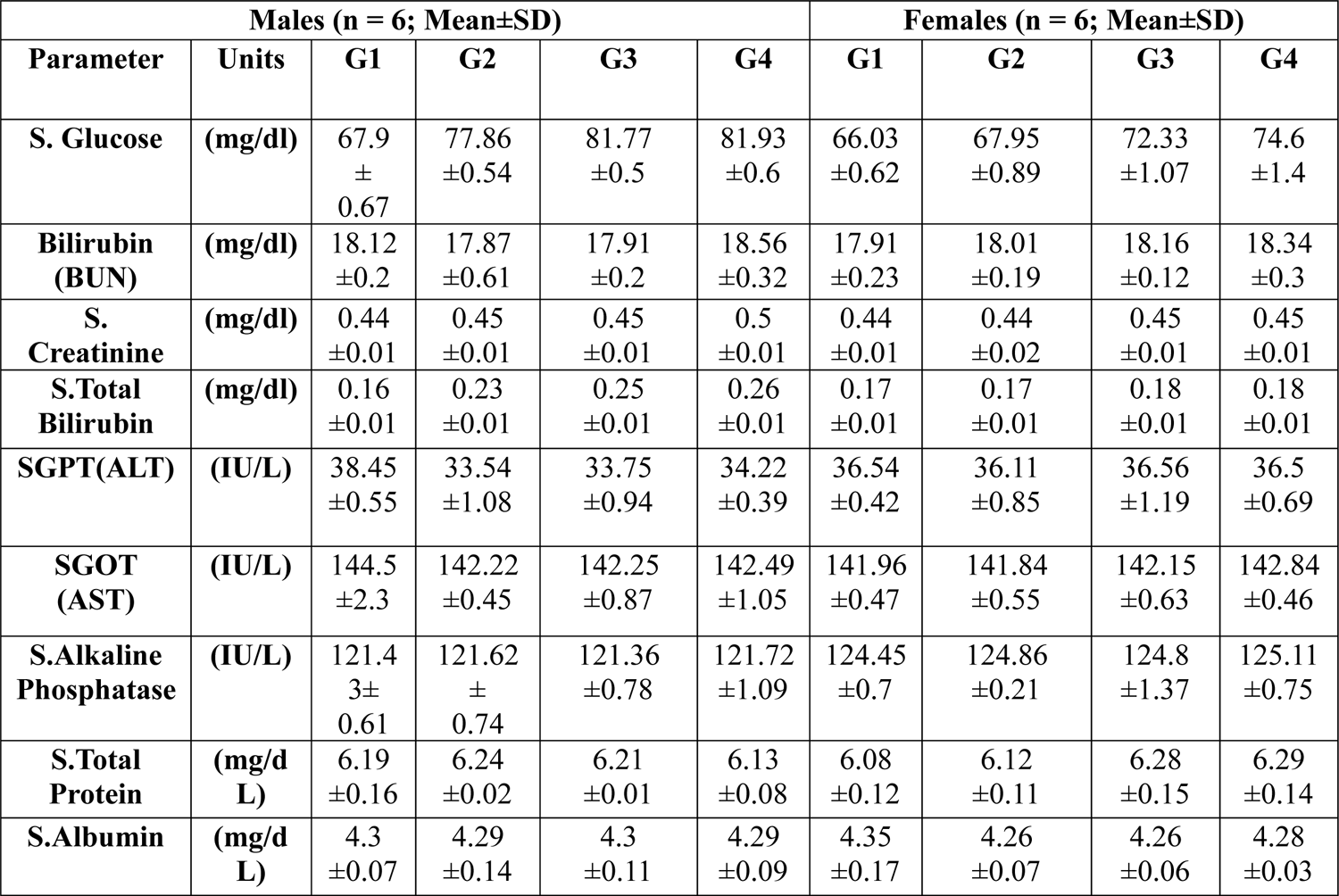
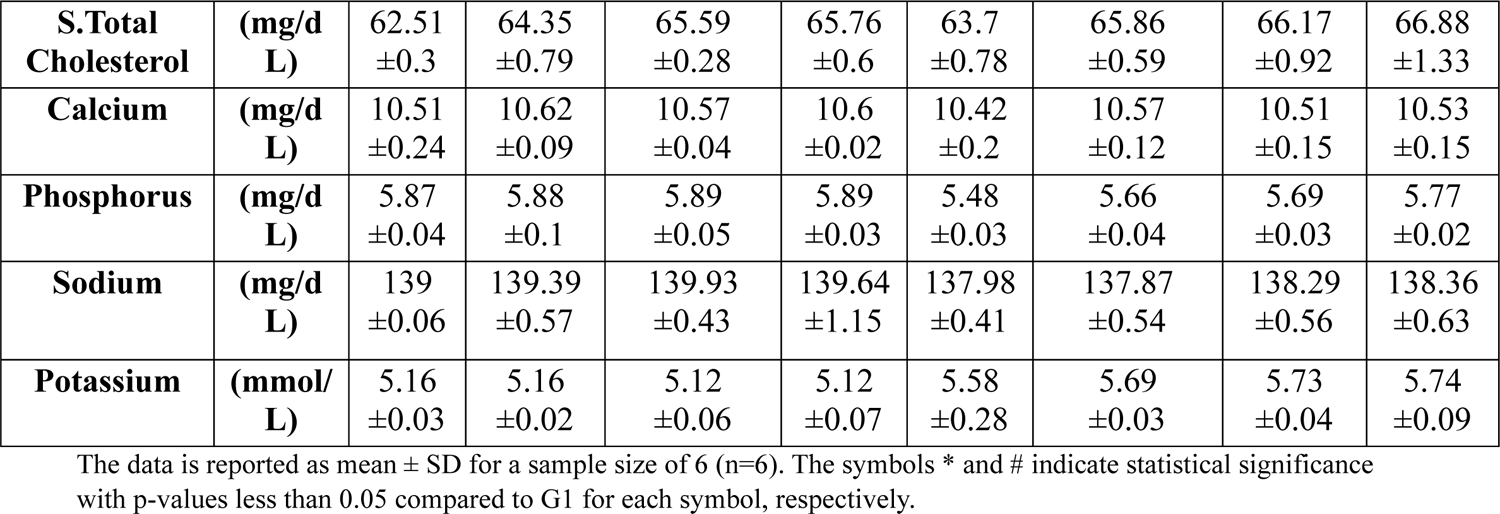
Summary of clinical chemistry parameters in Wistar Rats.

The analysis of haematological parameters, as depicted in Table 8, demonstrates that both the drug-treated and vehicle control groups exhibited haematological values within the normal range. These findings indicate that GlublocTM does not significantly impact the blood’s cellular or non-cellular components even at high dose levels. Hence, considering the aforementioned results, it can be concluded that Glubloc^TM^ did not induce any significant effects on hematological parameters throughout the 28-day repeated oral toxicity studies conducted on rats.

The findings presented in Table 9 indicate no significant alterations in the biochemical parameters. Most observed changes were non-significant and did not exhibit a dose-dependent pattern. Furthermore, these changes fall within the normal range, indicating that they cannot be categorized as pathological. Table 5 illustrates that the values of additional biochemical parameters remained within the normal range, even when higher doses of Glubloc^TM^ were administered. No significant alterations were observed in the biochemical parameters of the groups treated with the test drug. Based on these findings, it can be inferred that the test drug does not appear to induce any significant toxic effects during the 28-day repeated oral toxicity study. During the histopathological analysis of essential and reproductive organs at higher doses of Glubloc^TM^, no discernible changes were observed in the cellular structure of the brain, liver, heart, spleen, kidney, lung, pancreas, epididymis, testes, uterus, ovary, and adrenal gland when compared to the control group. The results suggest that the drug does not elicit any adverse effects on organs even at high dose levels.

The daily administration of Glubloc^TM^ at a dose of 414.16 mg/kg BW did not lead to notable changes in the vital organs of male and female experimental animals. Throughout the treatment period, there were no observable alterations in the gross morphology of the essential organs in the male and female rats subjected to high-dose administration. Table 10 provides a summary of the histological observations conducted on the vital organs of male animals, while Table 11 presents a summary of the histological observations on the vital organs of female animals.

**TABLE 10:**
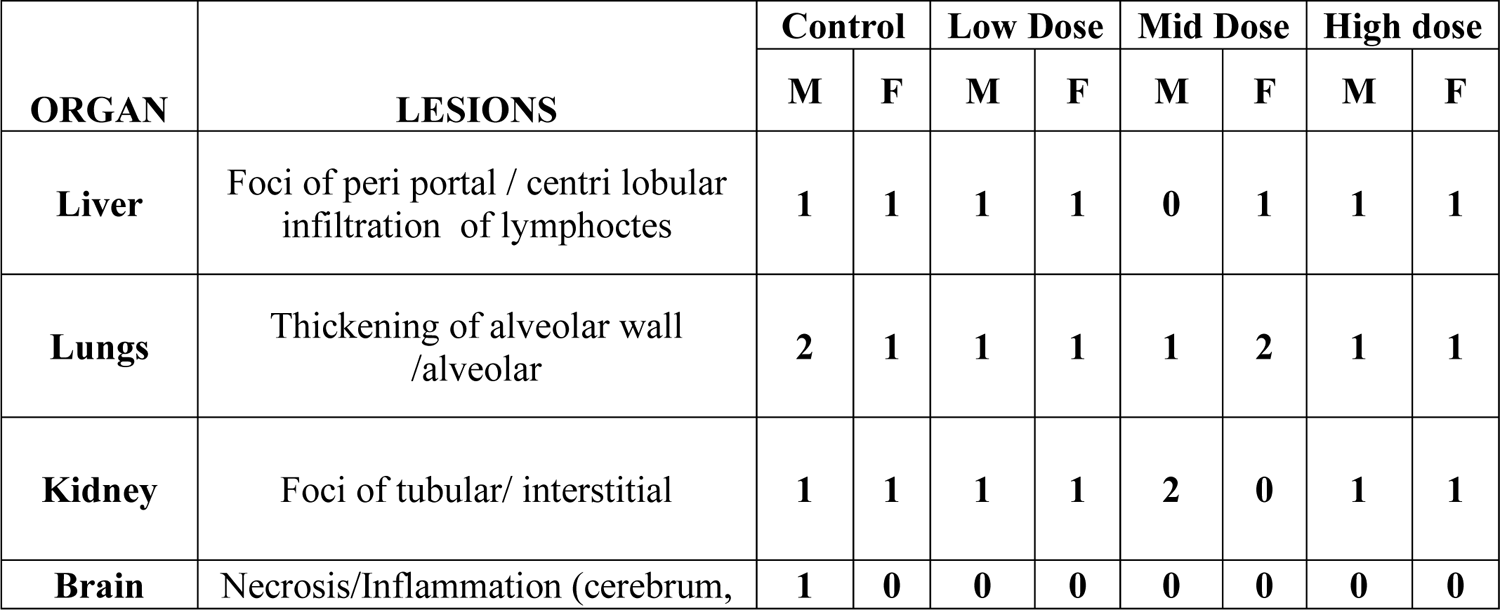
Summary of histopathological observation in Wistar Rats.

**TABLE 11:**
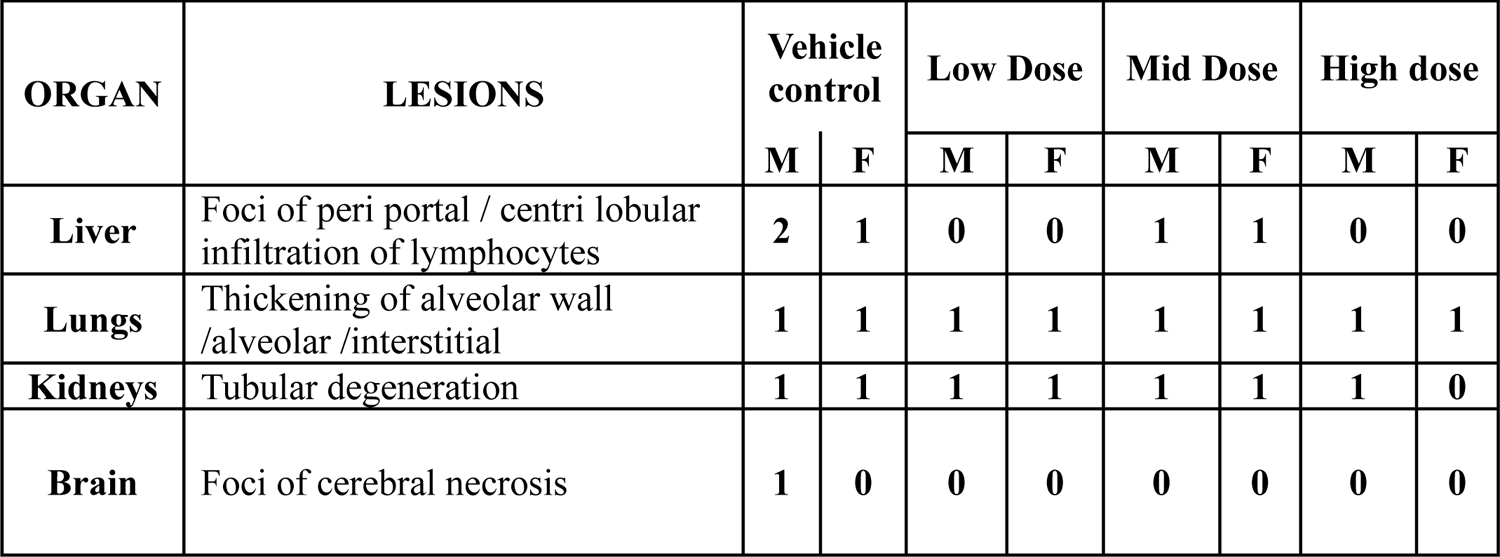
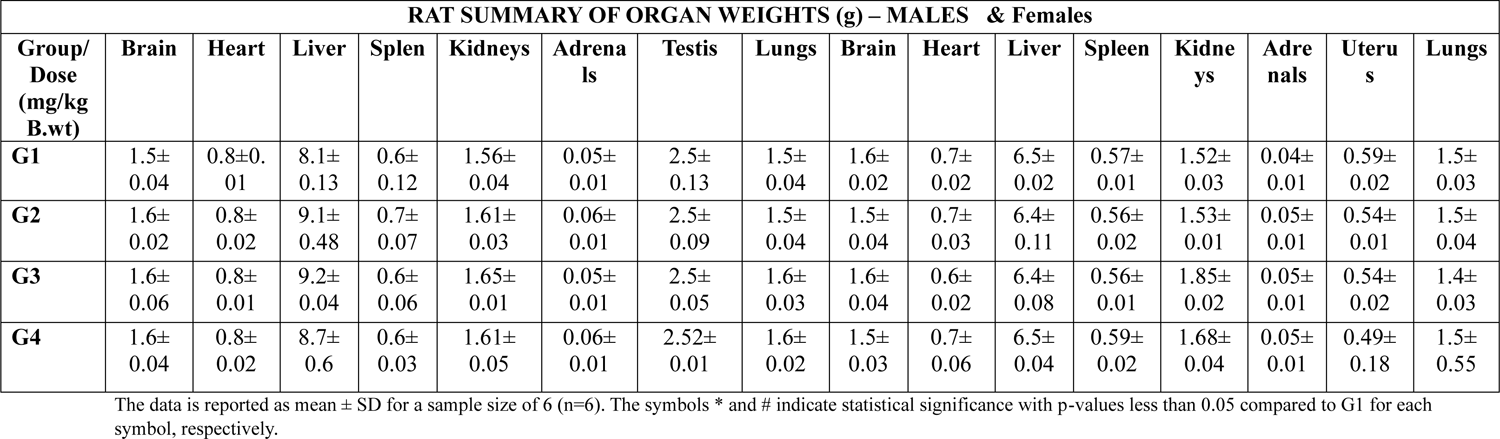
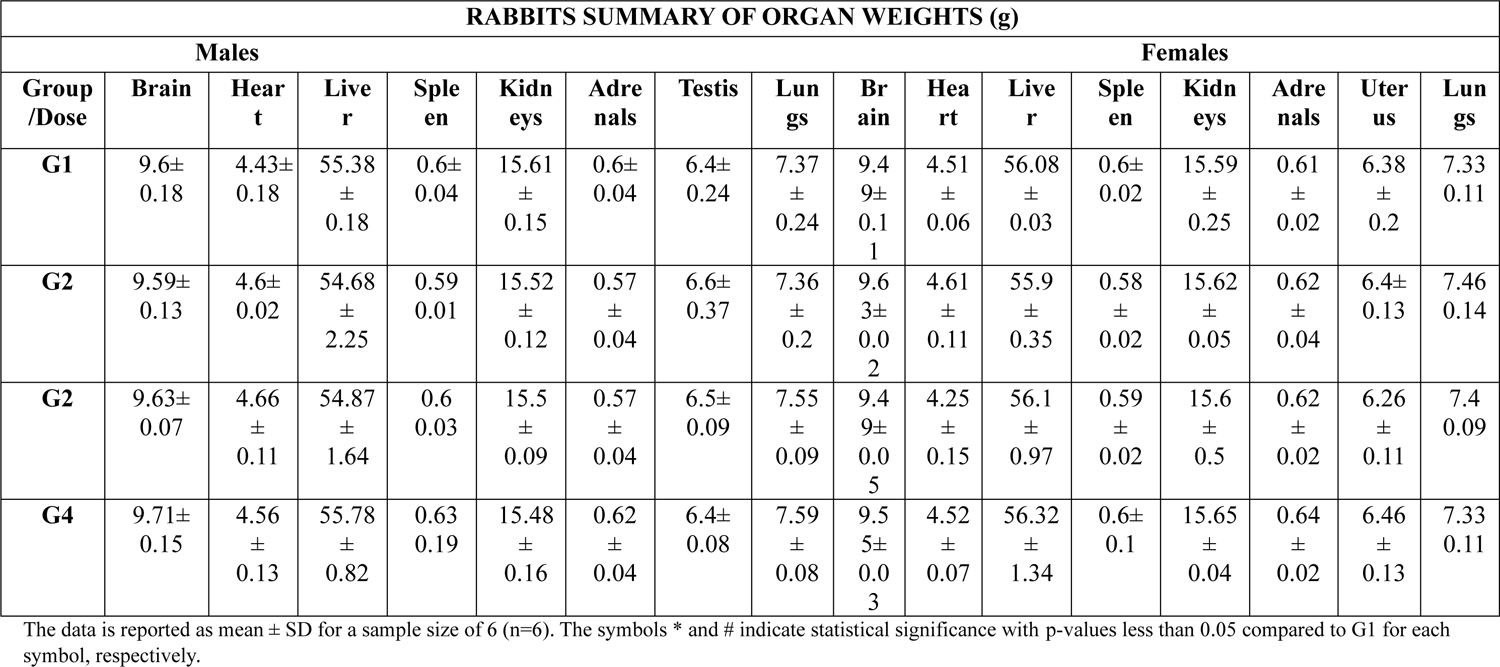
Summary of histopathological observation in rabbits.

## 4. Discussion

For centuries, both of these plants have held significant medicinal value in complementary and alternative medicine, each with unique properties. Due to their extensive historical use as both medicinal agents and food ingredients, these plant materials are generally regarded as safe for human consumption—the usage history of the traditional medicinal.

In accordance with the guidelines established by the OECD, we conducted toxicity evaluations of the Glubloc^TM^ via acute oral toxicity and subacute twenty-eight-day repeated dose oral toxicity studies in both rats and rabbits.

*Malus Domestica* and *Morus alba* (white mulberry) contains various bioactive components such as polyphenols, flavonoids, alkaloids, and imino sugars, which are believed to affect cellular pathways and molecular targets involved in glucose metabolism. Polyphenols and 1-deoxynojirimycin (DNJ) found in these plants can inhibit carbohydrate-digesting enzymes, which may result in lower postprandial glucose levels. In contrast, the modulation of glucose transporters and enhancement of insulin sensitivity can help improve glucose uptake and utilization by cells. Moreover, the anti-inflammatory and antioxidant properties of *Malus domestica* and *Morus alba* can potentially reduce oxidative stress and inflammation linked to insulin resistance and diabetes.

The initial step in our study was to establish that the oral LD_50_ of Glubloc^TM^ was the highest tested dose of 2000 mg/kg BW in Wistar rats and albino mice. The herbal composition exhibited no signs of acute toxicity, mortality, or abnormalities in 14 days of observation. Based on the OECD’s toxicological classification criteria, the findings indicate that the test item is a non-toxic composition and can be classified under the “no label” category.

In order to minimize the use of animals and decrease variability, the OECD 425 guideline limit test for oral acute toxicity study was exclusively performed on female rats, as they were generally found to be slightly more sensitive. The limit test is typically employed when prior knowledge suggests that the test substance is expected to have little or no toxicity. In the case of Glubloc^TM^, no signs of toxicity or mortality were observed in female rats at an oral dose of 2000 mg/kg, and there were no changes in behaviour or other observed parameters during the 14-day experimental period. This suggests that the oral LD_50_ of Glubloc^TM^ is much higher than the administered dose of 2000 mg/kg in the present study. In accordance with the UN Classification, substances that possess an oral LD_50_ more fabulous than 2000mg/kg are regarded as having low hazard potential and are classified as UN 6.1 PG III.

In the subacute toxicity study conducted for twenty-eight days, rats that were given the highest dose of 414.16 mg/kg/day of Glubloc^TM^ did not exhibit any observable signs of toxicity. No animal in either the primary died during the study. Furthermore, no treatment-related changes were observed in body weight, cumulative feed consumption, gross anatomy, or histopathology in the animals.

Throughout the current subacute study, the animals belonging to the treatment groups showed a gradual increase in their body weights. There was no correlation observed between the amount of feed consumed by these groups and their body weight data. Additionally, the total feed consumption in these groups did not show any significant difference compared to their respective controls. Taken together, these observations indicate that the experimental rats’ natural growth and metabolism remained unaffected by the oral administration of Glubloc^TM^.

The hematopoietic system is a highly vulnerable target of toxic substances and serves as a critical indicator of both normal and abnormal physiological conditions in humans and animals. The administration of the Glubloc^TM^ for 28 days did not change the hematological parameters of the treated group compared to the control group. This indicates that the extract did not have a toxic effect on the production and circulation of red blood cells and platelets, which are highly sensitive to harmful substances. Nevertheless, these hematology and clinical chemistry values fall within the normal physiological ranges associated with the age and gender of rats and rabbits. These changes are not correlated with either the dosage or the duration of the treatment.

Furthermore, the extract did not cause any notable changes in the majority of the biochemical parameters tested. The absence of significant modifications in the levels of liver and kidney function markers such as ALT, AST, alkaline phosphatase, glucose, and creatinine indicates that the administration of the extract over a sub-chronic period did not adversely affect the liver, kidneys or metabolism of the rats.

The kidneys eliminate urea and creatinine from the body through urine. Elevated levels of these metabolites in the bloodstream indicate a potential impairment in kidney function, as they may indicate insufficient clearance from the body. The current study observed that the oral administration of Glubloc^TM^ did not influence the serum levels of urea and creatinine in both male and female rats and rabbits. This indicates the absence of any detrimental effects of the treatment on the kidneys of the experimental animals. Overall, the observations on the hematobiochemical parameters suggest that the maximum tested dose of the herbal blend (414.16 mg/kg/day in rats and 207.08 mg/kg in rabbits) did not result in pathological changes in the treated animal’s vital organs.

Furthermore, both the macroscopic and microscopic evaluations of the vital organs in rats and rabbits revealed no treatment-related changes. These observations provide further support that the oral administration of this Glubloc^TM^ did not result in any systemic toxicity in male and female rats.

Collectively, the outcomes of the twenty-eight-day repeated oral toxicity investigation confirm that the NOAEL of Glubloc^TM^ in male and female rats is 414.16 mg/kg body weight per day and rabbits is 207.08 mg/kg body weight which was the highest administered dose in the study. The histopathological evaluations of the vital and reproductive organs at the higher dose level indicated no changes in the cellular structure of the brain, pituitary, liver, heart, thymus, spleen, kidney, lung, uterus, and ovary. And adrenal gland as compared to the control and vehicle control group. Certain medications and drugs can cause liver damage, leading to foci of periportal inflammation. One animal showed foci of periportal/centrilobular infiltration in both male and female groups. However, the architecture of the liver in mid-dose male rats and low/high doses of rabbits remained unchanged. The results may not be considered significant since the observed changes were present in both the control and treatment groups.

The control group rats and rabbits exhibited changes in the thickening of the alveolar wall in both sexes. Among the animals, two males from the control group and two from the mid-dose showed the thickening of the alveolar wall. All the rats have shown pathological changes in the kidneys by foci of tubular/ interstitial. 2 animals from the mid-dose have reflected changes in the kidney architecture. Exposure to toxins, such as heavy metals or solvents, can cause damage to the kidneys. The observed pathological changes were not considered significant, as they were also present in the control group. One control group, male rat and rabbits have shown that foci of cerebral necrosis may not be considered significant as they were the control group.

## Conclusion

The present study’s findings suggest that oral administration of Glubloc^TM^ at a single dose of 2000 mg/kg did not manifest any observable signs or symptoms of acute toxicity in mice and rats. Therefore, it can be concluded that Glubloc^TM^ has a low potential for health hazards. The result of a repeated dose of 28-day oral toxicity concluded that the test drug was safe without any toxic effects at the therapeutic level of 414.16 mg/kg BW in rats and 207.08 mg/kg BW in rabbits.

---

**Fig:**
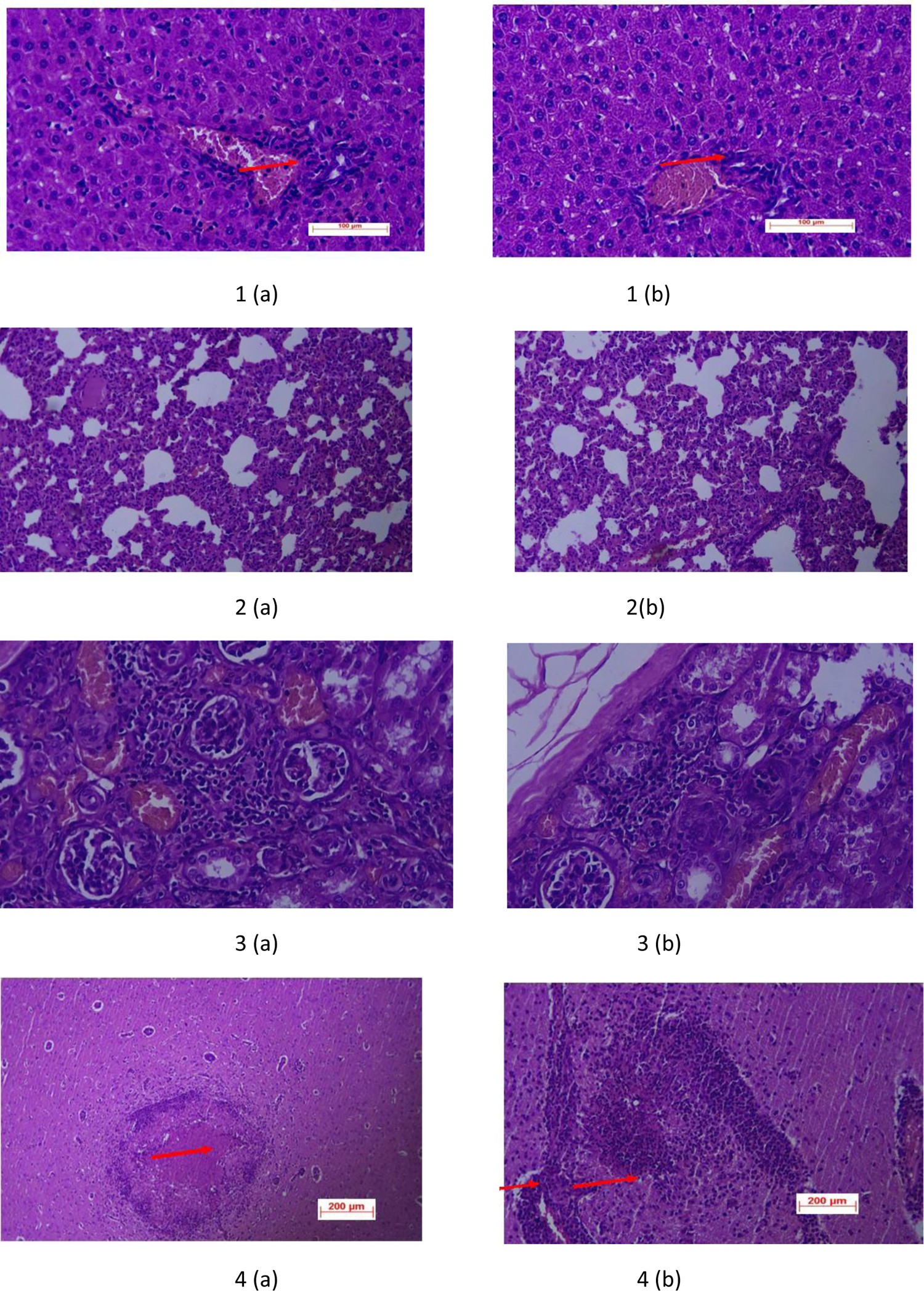

Histopathology of Wistar rat organs of control and treatment groups 1a, 1b (liver)- mild periportal infiltration of inflammatory cells (lymphocytes) were noticed, red arrow both in the control and low, mid and high doses 2a, 2b(lungs)- Moderate alveolar/ interstitial wall thickening or connective tissue proliferation was observed – red arrow both in the control and low, mid and high doses 3a, 3b Severe tubular nephritis / Nephropathy: Multifocal tubular/interstitial inflammation [red arrow] with infiltration of lymphocytes was observed both in the control and low, mid and high doses 4a, 4b- severe multifocal necrosis and inflammation noticed in the cerebral cortex region of the brain in the control group.

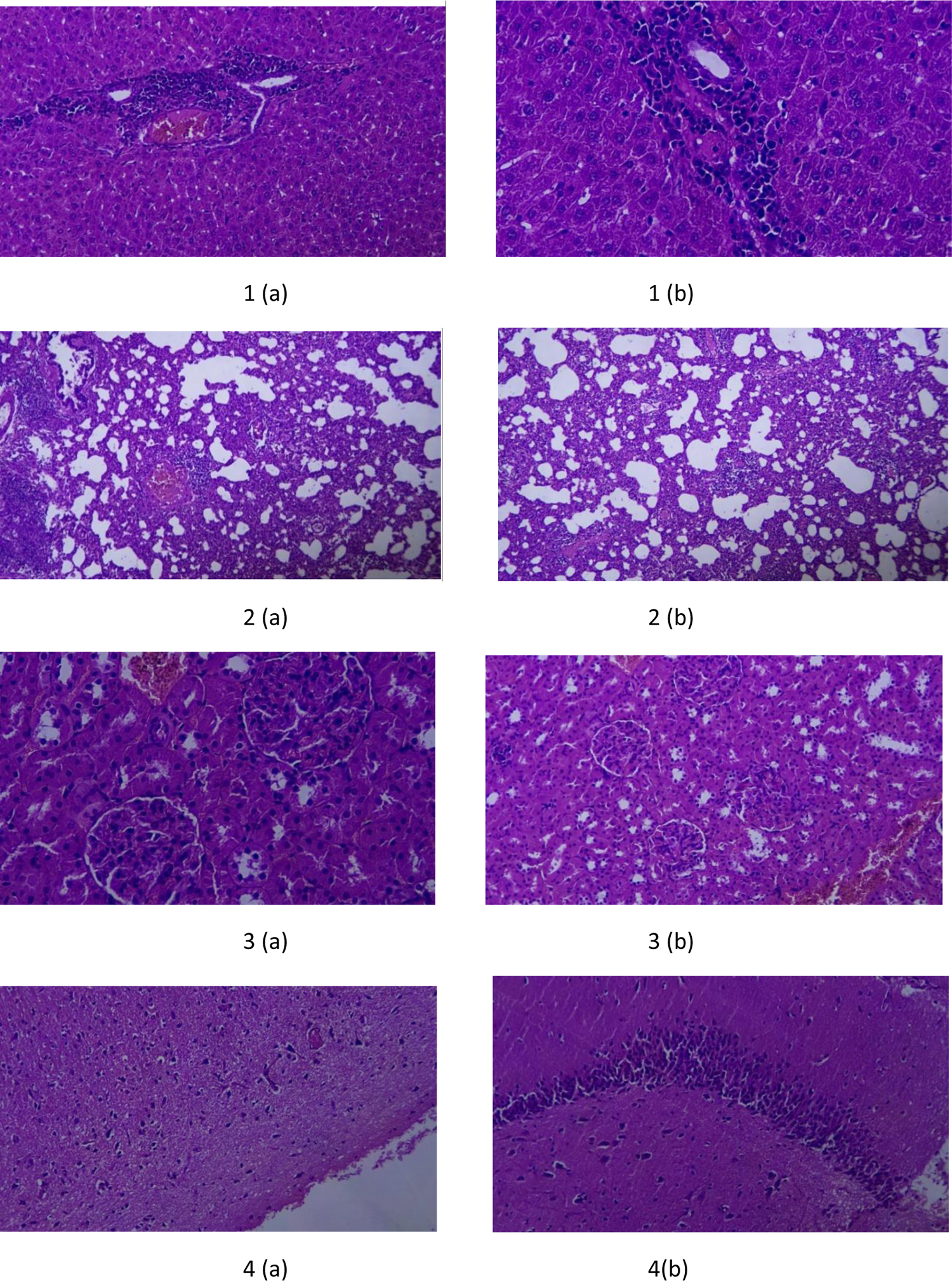

Histopathology of rabbit organs of control and treatment groups 1a, 1b (liver)- mild periportal infiltration of inflammatory cells (lymphocytes) were noticed, both in the control and mid doses 2a, 2b(lungs)- Moderate alveolar/ interstitial wall thickening or connective tissue proliferation was observed –both in the power and low, mid and high doses 3a, 3b Severe tubular nephritis / Nephropathy: Multifocal tubular/interstitial inflammation with infiltration of lymphocytes was observed both in control, low, mid and high doses 4a, 4b- severe multifocal necrosis and inflammation noticed in the cerebral cortex region of the brain in the control group.

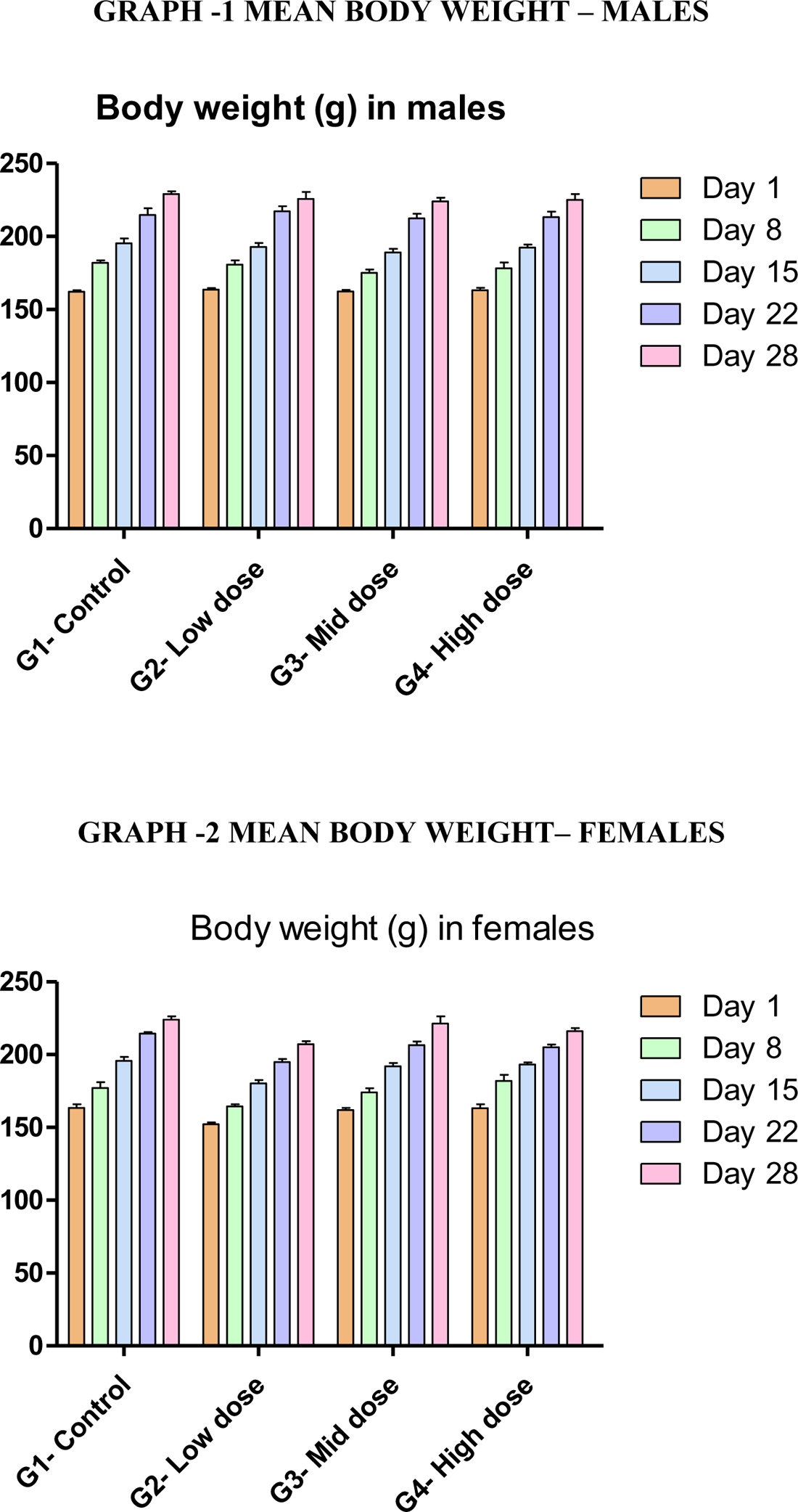

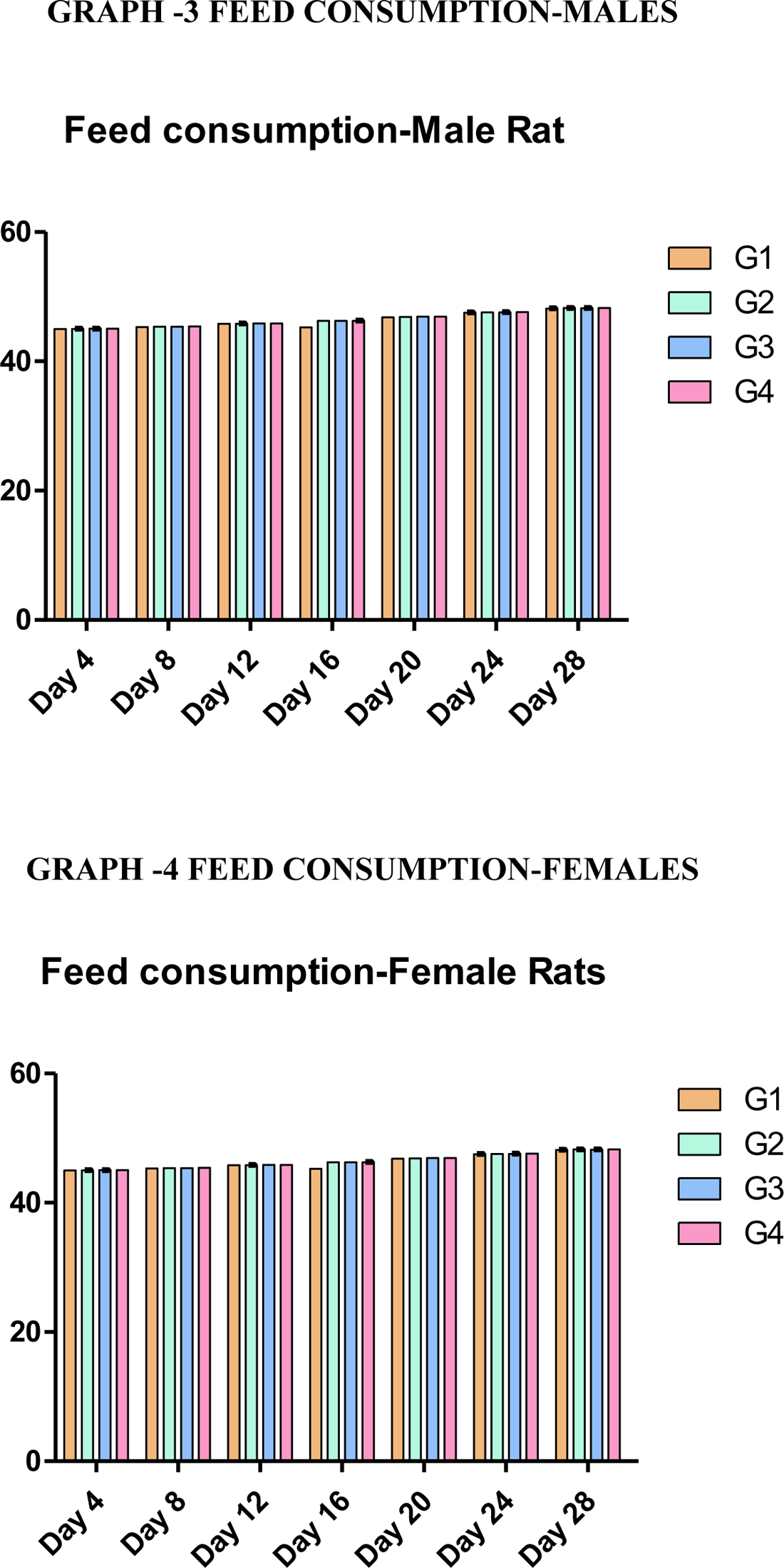

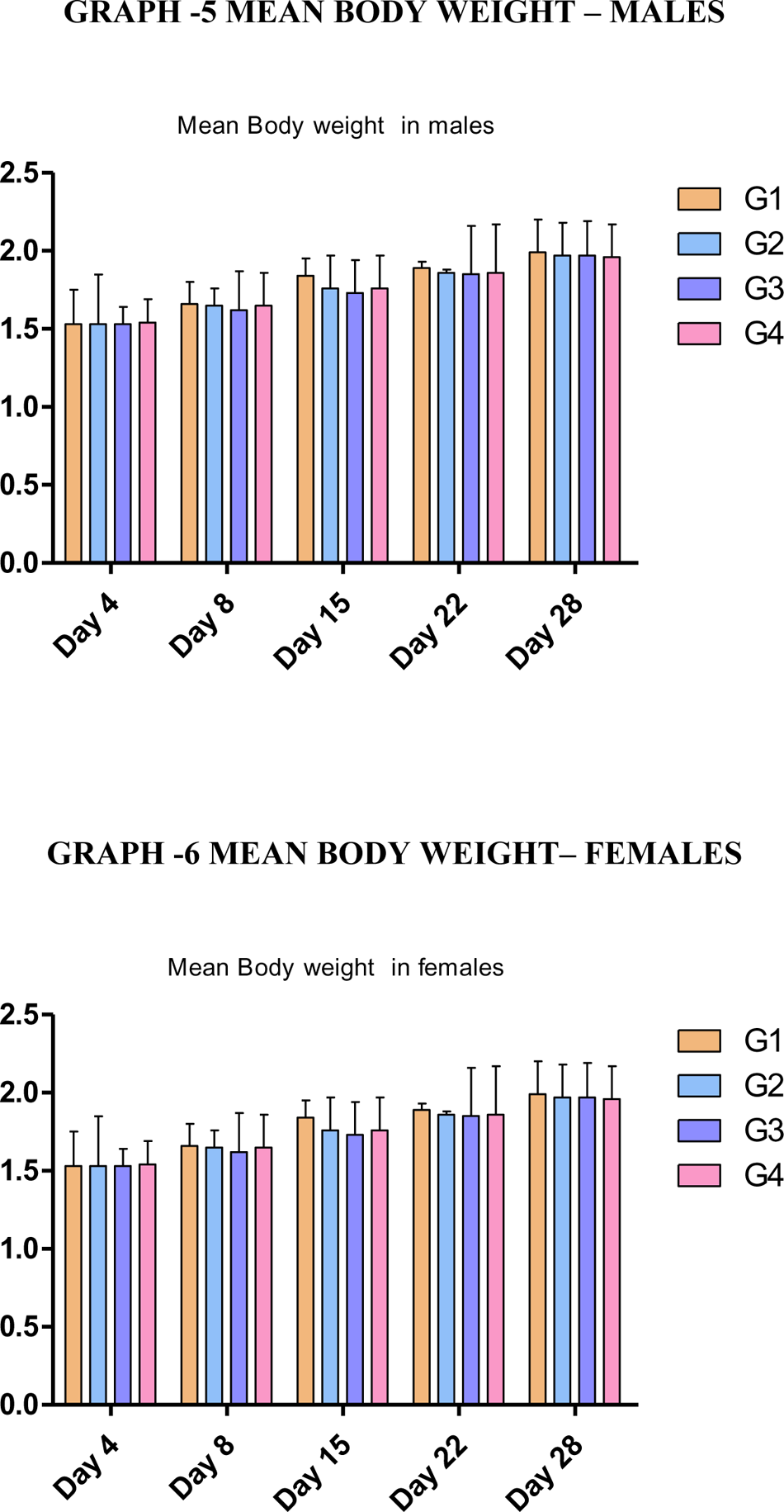

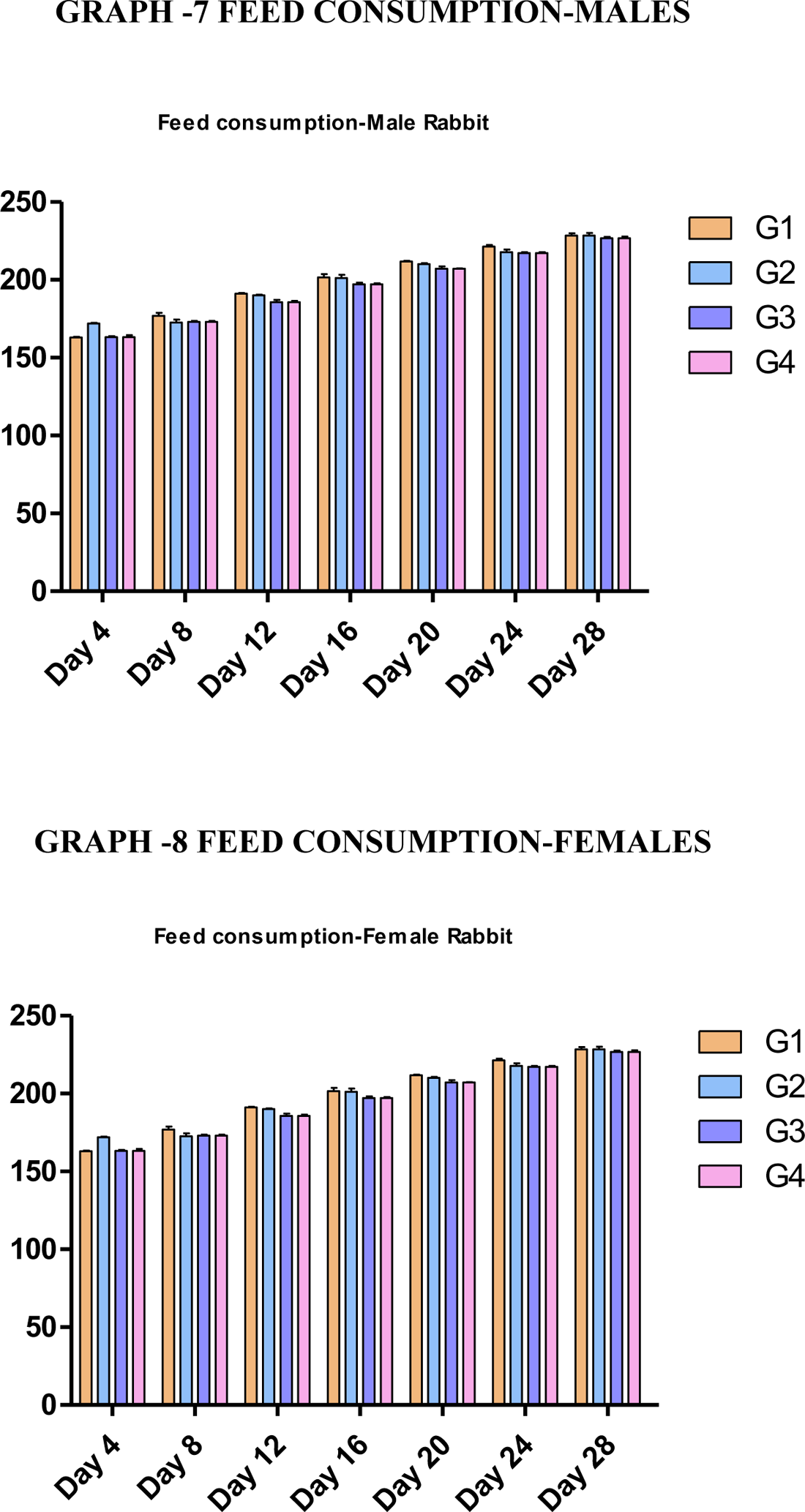

